# The NaaS Methodology applied to modeling chemotherapy-induced peripheral neuropathy with human hiPSC neurons

**DOI:** 10.1101/2025.09.25.678500

**Authors:** Thomas Bessy, Théo Lambert, Louise Dubuisson, Aurélie Batut, Audrey Azema, Camille Baquerre, Alexandre Ponomarenko, Serge Roux, Najate Ftaich, Thibault Honegger

## Abstract

The Neuron-as-a-Sensor (NaaS) methodology is a human-relevant platform designed to detect compound-induced effects by capturing functional changes in neuronal activity. This is achieved by integrating hiPSC-derived neuronal cultures, compartmentalized MEA microfluidic devices, a detailed electrophysiology paradigm and a standardized analysis pipeline. Applied to chemotherapy-induced peripheral neuropathy (CIPN), NaaS leverages electrophysiological profiling to capture alterations in neuronal excitability beyond cytotoxicity. Using paclitaxel and oxaliplatin as reference compounds, we demonstrated drug-specific, time-dependent changes in spontaneous and thermally evoked activity that align with their distinct clinical neuropathic phenotypes. Dimensionality reduction of electrophysiological metrics enabled construction of a functional discrimination map, allowing robust separation of compound signatures from vehicle controls. These findings highlight the ability of NaaS to model clinically relevant neurotoxic effects in a scalable manner, supporting its application in both adverse effect prediction and therapeutic screening.

## INTRODUCTION

Drug development continues to face significant challenges, with high attrition rates in the transition from preclinical to clinical phases. A major contributor to this attrition is the limited predictive power of current preclinical models, which often fall short in forecasting clinical outcomes related to both efficacy and safety ^1^. This disconnect leads to costly late-stage failures and highlights a critical need for more human-relevant and mechanistically informative systems early in the discovery pipeline.

Rodent models remain a cornerstone of preclinical testing due to their well-characterized biology and utility in mechanistic research. However, growing evidence has revealed substantial species-specific differences, particularly at the molecular and transcriptomic levels ^2,3^. These differences undermine their translational relevance, especially in complex phenotypes such as pain or neurotoxicity, where clinical outcomes are poorly predicted ^4^. Additionally, animal studies pose scalability challenges and raise ethical concerns, further limiting their suitability for high-throughput screening.

In response, human cell-based systems have gained traction. Two-dimensional (2D) cultures of immortalized or primary human cells provide a more ethically acceptable and human-relevant alternative. Among all cell-based assays, neurons derived from human induced pluripotent stem cells (hiPSCs) represent an interesting choice for a model. These cells combine human origin with the ability to recapitulate key aspects of native neuronal function. Importantly, neurons are inherently electrically active and responsive to pharmacological and toxicological stimuli, which allows their functional status to be monitored dynamically over time. Multi-electrode arrays (MEA) in particular, facilitate real-time, label-free, and non-destructive tracking of neuronal network activity, providing a rich dataset for assessing both therapeutic action^5^ and adverse responses ^3^. However, 2D culture models often lack the structural and functional complexity needed to capture physiological responses.

To address these limitations, New Approach Methodologies (NAMs), including organoids and organ-on-chip (OOC) systems, have emerged to address these gaps by recapitulating tissue architecture and cell-cell interactions. While promising, these models frequently suffer from low throughput, limited reproducibility, and a lack of standardized protocols, all of which hinder their widespread adoption ^6^.

In this context, we developed the Neuron-as-a-Sensor (NaaS) methodology, a comprehensive hardware, software and biology platform that integrates hiPSC-derived neuronal cultures with NETRI’s 96-well format microfluidic devices equipped with embedded multi-electrode arrays (MEAs). The methodology includes a standardized protocol for longitudinal electrophysiological monitoring and a dedicated data analysis pipeline. The aim is to provide a robust and scalable methodology capable of comparatively assessing compound effects in context-specific biological settings, including the innervation of targeted 2D/3D cultures. The NaaS methodology is designed not only to detect overt toxicity but also to capture nuanced, functional alterations in neural activity over time.

To demonstrate the relevance and capabilities of the NaaS platform, we applied it to a high-need clinical context: chemotherapy-induced peripheral neuropathy (CIPN). CIPN is a frequent and debilitating side effect of many cancer treatments, characterized by sensory abnormalities and chronic pain. Despite its prevalence, CIPN remains poorly modeled and under-addressed in drug development. Current preclinical assays, either rodent or cell-based, fail at predicting compounds efficacy with regards to neuropathies ^7,8^. Because CIPN reflects a complex, functional disruption of peripheral neurons—rather than a simple loss of viability—it serves as an ideal test case for a system designed to capture longitudinal neural dysfunction in a human-relevant context.

In summary, by leveraging the electrophysiological properties of hiPSC-derived neurons and integrating them with robust hardware and analytics, the NaaS platform aims at addressing key gaps in the potency of existing models for predicting adverse effects or efficacy in a range of biomedical contexts.

## RESULTS

The aim of the NaaS methodology is to leverage the discriminative power of comprehensive MEA data to predict the properties of novel compounds. To achieve this outcome, we propose to describe the electrophysiological activity of a neuronal culture in comparison with its alteration when exposed to a specific compound. To characterize this difference, the methodology has to be comprised of: a set of well-documented reference compounds to map the state of the clinical context, a cell culture that is relevant to the biological question, a NETRI compartmentalized MEA device for a physiologically relevant cell organization (**Figure 1.a, left**). An experimental methodology with a set of descriptive recordings will define the electrophysiological signature of the chosen reference compounds, which corresponds to the shift of activity with regards to untreated neurons. These deltas are then used to build a reference map (**Figure 1.a, right**), in which an unknown compound – tested under the same descriptive recordings -can be positioned to infer its properties on neurons (**Figure 1.b**).

**Figure 1.**
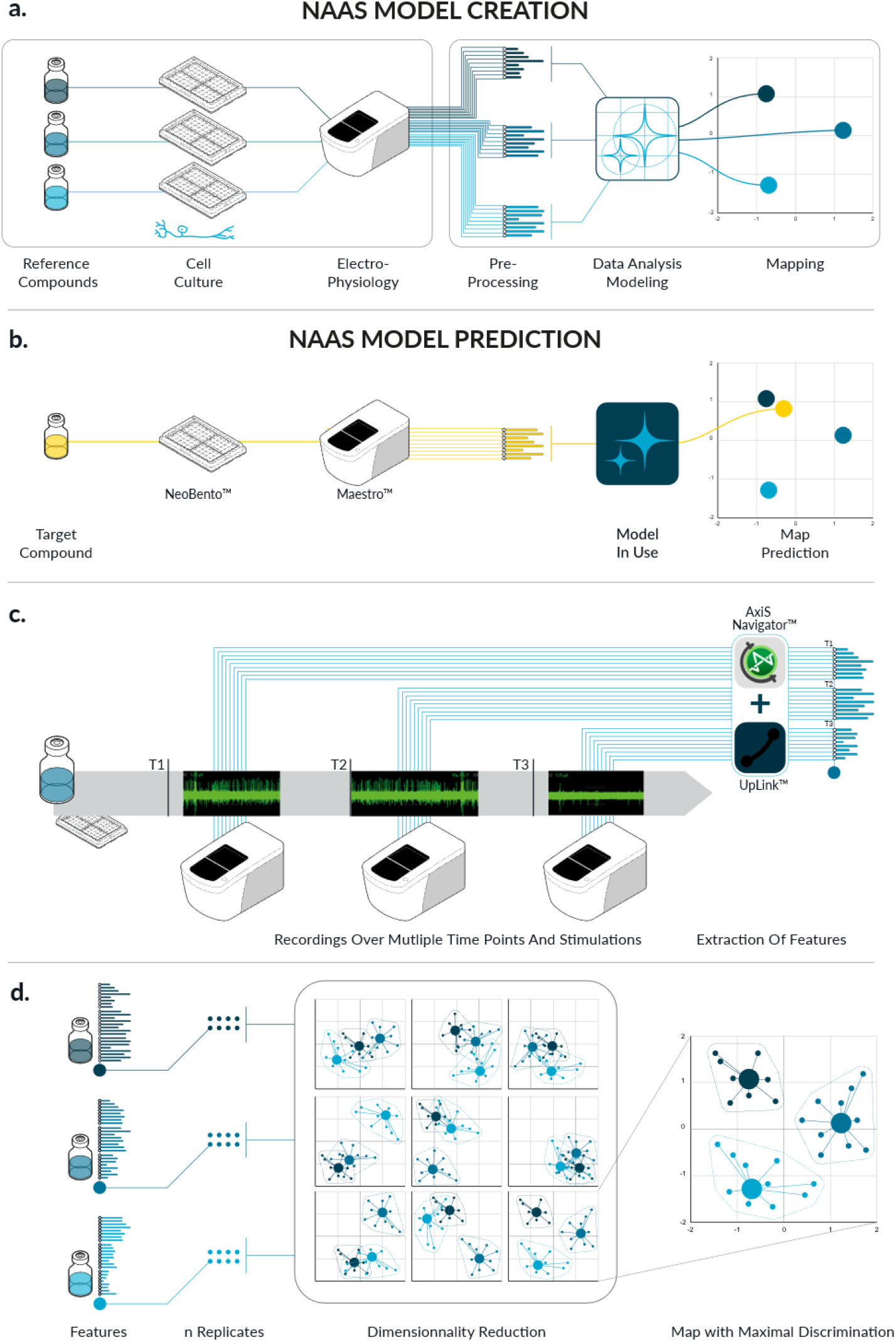
Methodology of the Neuron-as-a-Sensor (NaaS) Platform. **a**. *Left:* Reference compounds are applied to neuronal cell cultures grown on NETRI compartmentalized MEA devices. Electrophysiological responses are recorded using Maestro PRO MEA system. *Right:* Extracted electrophysiological features are processed using a dimensionality reduction algorithm to map compound effects within a relevant low-dimensional subspace. **b**. To assess an unknown compound, the same experimental protocol is applied. Electrophysiological features are extracted and projected into the reference space to infer the compound’s functional profile. **c**. Electrophysiological activity is recorded at multiple time points and under different conditions. Signals are processed using AxisNavigator (Axion Biosystems) and NETRI UpLink to extract a wide range of electrophysiological metrics describing the culture’s functional state. **d**. Features obtained across time points and conditions are subjected to dimensionality reduction. The axes offering the highest discriminatory power are retained to serve as the basis for comparison with novel compounds.

As part of the experiment plan, the goal is to generate a series of descriptive functional states of the neuronal culture subjected to a compound through electrophysiology. To achieve this, cells will be recorded over multiple time points and stimulations (**Figure 1.c**). Each recording helps describe the neuron firing pattern and its excitability through time, for an exhaustive description of its biological state. Signals are then processed with tailored software to obtain a large set of descriptive features, comprised of various metrics in the mentioned functional states.

Once these features are obtained for reference compounds, they are processed through a dimensionality reduction model to obtain low-dimensional subspaces in which references are well separated (**Figure 1.d**). The same procedure is applied to test a new compound with respect to the reference compounds.

### Establishing the Neuron-as-a-Sensor Platform for CIPN Applications

To evaluate the relevance and utility of the NaaS methodology in modeling chemotherapy-induced peripheral neuropathy (CIPN), we established an experimental framework combining compound selection, biologically relevant cell types, microphysiological hardware, and a functional assay paradigm that captures clinically relevant electrophysiological phenotypes.

We selected two chemotherapeutic agents—paclitaxel and oxaliplatin—based on their well-documented yet distinct clinical association with CIPN^9^. Paclitaxel, a taxane, disrupts microtubule dynamics by stabilizing tubulin polymers, ultimately impairing axonal transport. It is commonly associated with chronic sensory neuropathy and thermal hyperalgesia in patients. Oxaliplatin, a platinum-based agent, exerts its anticancer effects via DNA crosslinking but is also known to induce acute neuropathy characterized by cold hypersensitivity.

To model peripheral sensory function, we employed a co-culture of hiPSC-derived sensory neurons and glial cells (astrocytes). hiPSC derived sensory neurons have previously been described in their potency to model neurotoxic effect of CIPN inducing drugs^10–12^. Moreover, co-culturing sensory neurons with glial cells provides a more physiologically relevant microenvironment, as glia support neuronal survival, maturation, and stable electrophysiological activity ^13,14^. Cultures were seeded into NETRI’s DuaLink microfluidic chips with integrated multi-electrode arrays (MEAs). These chips were selected for their suitability in high-throughput electrophysiological studies and their compartmentalized architecture, which separates neuronal somas from distal processes via microchannels. This configuration enables physiological organization and signal amplification in microchannel electrodes^15^.

To model functional disruptions while avoiding cytotoxic effects, co-cultures were treated with paclitaxel at 1 µM and oxaliplatin at 10 µM. These concentrations are the order of magnitude of Cmax plasma levels for paclitaxel^16^ and of the accumulated concentration of platin in the tissues for oxaliplatin^17^. They were further validated in our system using LDH release assay after 48 hours of exposure. Compounds were administered to all compartments of the microfluidic chip, simulating systemic drug exposure as occurs *in vivo*. Cell viability was maintained above 95% under both conditions, confirming sub-cytotoxic exposure (data not shown).

To probe for early electrophysiological changes relevant to CIPN, we implemented an experimental schedule incorporating longitudinal recordings and thermal stimulations. Recordings were performed at baseline (prior to treatment), and at 24- and 48-hour post-treatment. Recording the initial state of the culture allows for quality control by excluding devices with insufficient activity and enabled the use of an algorithm to assign devices to treatment groups with homogeneous baseline activity, in accordance with best practices for MEA electrophysiology studies^18^. At each time point, spontaneous and thermally evoked activity were assessed at 42°C. This temperature ramp was chosen to mimic clinical thermal hyperalgesia, a common symptom in CIPN patients, and to functionally challenge the system’s sensory responsiveness.

### Biological model development and characterization

To establish a physiologically relevant model for CIPN, hiPSC derived sensory neurons and cortical astrocyte were seeded in one of the compartment of NETRI DuaLink MEA microfluidic chip. A 2:1 ratio of 40,000 sensory neurons to 20,000 astrocytes was seeded into the somatic (left) compartment. Over time, axons progressively grew through the microchannels into the terminal endings compartment (right).

In preliminary experiments, immunofluorescence staining was performed on sensory neurons monocultures to confirm neuronal differentiation in the devices. The presence of key nociceptor differentiation markers expression, including TRPV1 and of the voltage-gated sodium channels Nav1.7 and Nav1.8, were confirmed (data not shown).

In the co-culture condition, further immunofluorescence analyses confirmed the presence and spatial organization of both cell types. Sensory neurons were stained with βIII-tubulin and MAP2, while astrocytes were identified via GFAP.

Sensory neurons were evenly distributed throughout the somatic compartment and astrocytes were located mainly around sensory neurons clusters. Neuronal extensions extended through microchannels and covered the entire distal channel. (**Figure 2.b**.)

**Figure 2.**
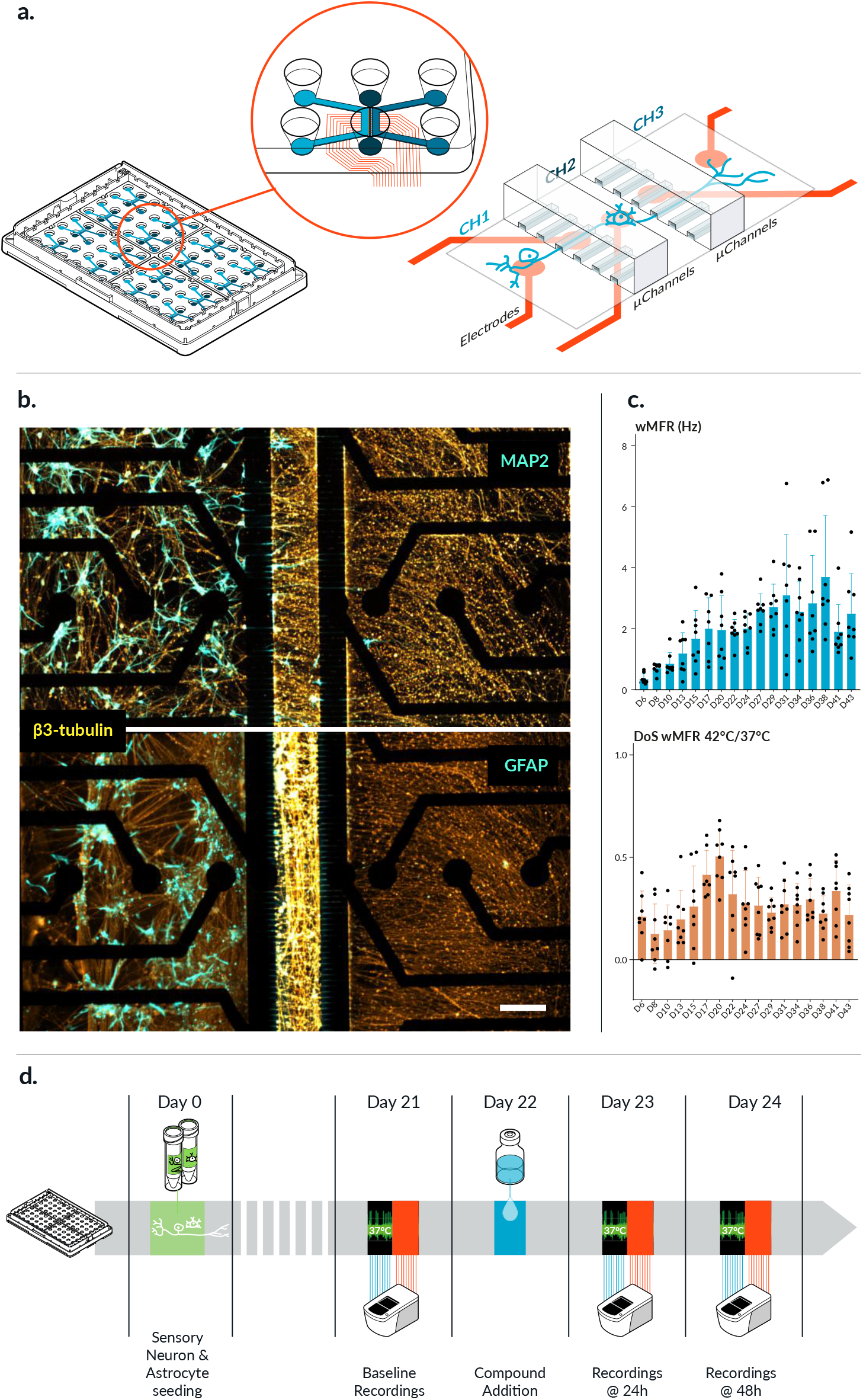
Development of the NaaS Biological Model for CIPN Applications. **a**. *Left:* Real-scale image of a NETRI DuaLink plate and device. *Right:* Schematic representation of a sensory neuron–astrocyte co-culture within a microfluidic chip. **b**. Representative immunofluorescence image of a sensory neuron–astrocyte co-culture at DIV 24, showing β3-tubulin (orange), MAP2 and GFAP (cyan) staining. Scale bar = 200 µm. **c**. Electrophysiological recordings of co-cultures were performed three times per week at 37°C and 42°C. *Left:* Weighted Mean Firing Rate (wMFR) over time. *Right:* Difference of Signal (DoS) between wMFR at 37°C and 42°C. **d**. Experimental paradigm diagram showing the assay timeline with scheduled recordings at 37°C (green) and 42°C (red).

To characterize co-cultures’ electrophysiological maturation, recordings were performed three times per week for 43 days. Basal activity was detected as early as D6 and reached a plateau from D17 to D24. Activity rose again from D24 but with increased dispersion of points until D38. Finally, activity decreased from D41 onwards. Temperature response to 42°C exposure was also recorded at each time point. The cells responded to temperature with a significant increase in firing rate compared to baseline (37°C) as early as D6 (**Figure 2.c.)**. The differential thermal response became more pronounced at D17. After D24, the temperature response decreased but remained positive and stable until the end of the culture. This reaction to temperature increase confirms the nociceptive identity of the sensory neurons.

Based on the combined analysis of spontaneous and thermally evoked activity, the optimal assay window was determined to be between D21 and D24. During this time slot, cultures exhibited both a stable basal activity and response to temperature stimulation.

### Data acquisition and preprocessing

All recordings were made using a Axion Biosystems’ Maestro PRO piloted by the AxisNavigator software. The preprocessing pipeline is presented in **Figure 3.a**. The raw data consists of metadata regarding the acquisition and data stored in a raw file. This file is processed through the NETRI UpLink software, which, along with AxisNavigator, transforms the raw electrical traces recorded by the electrodes in a set of metrics. These metrics are calculated at the electrode level through AxisNavigator, while UpLink addresses the compartmentalization matching the architecture of the microfluidic device. Importantly, in this study, only the data coming from electrodes located in the micro-channels were further processed. Indeed, the high resistance generated by the small structure ensures a high signal to noise ratio of axonal extracellular action potentials, as previously described in the literature^15,19,20^. Metrics were normalized with respect to the reference state (**Figure 3.b**), which represents a relative evolution rather than an absolute value. Together with the microfluidic devices that standardize the culture and the guidance of axons atop electrodes in microchannels, these experimental choices help mitigate the inherent variability of MEA^18^. The list of all considered metrics is presented in the Materials and Method section.

**Figure 3.**
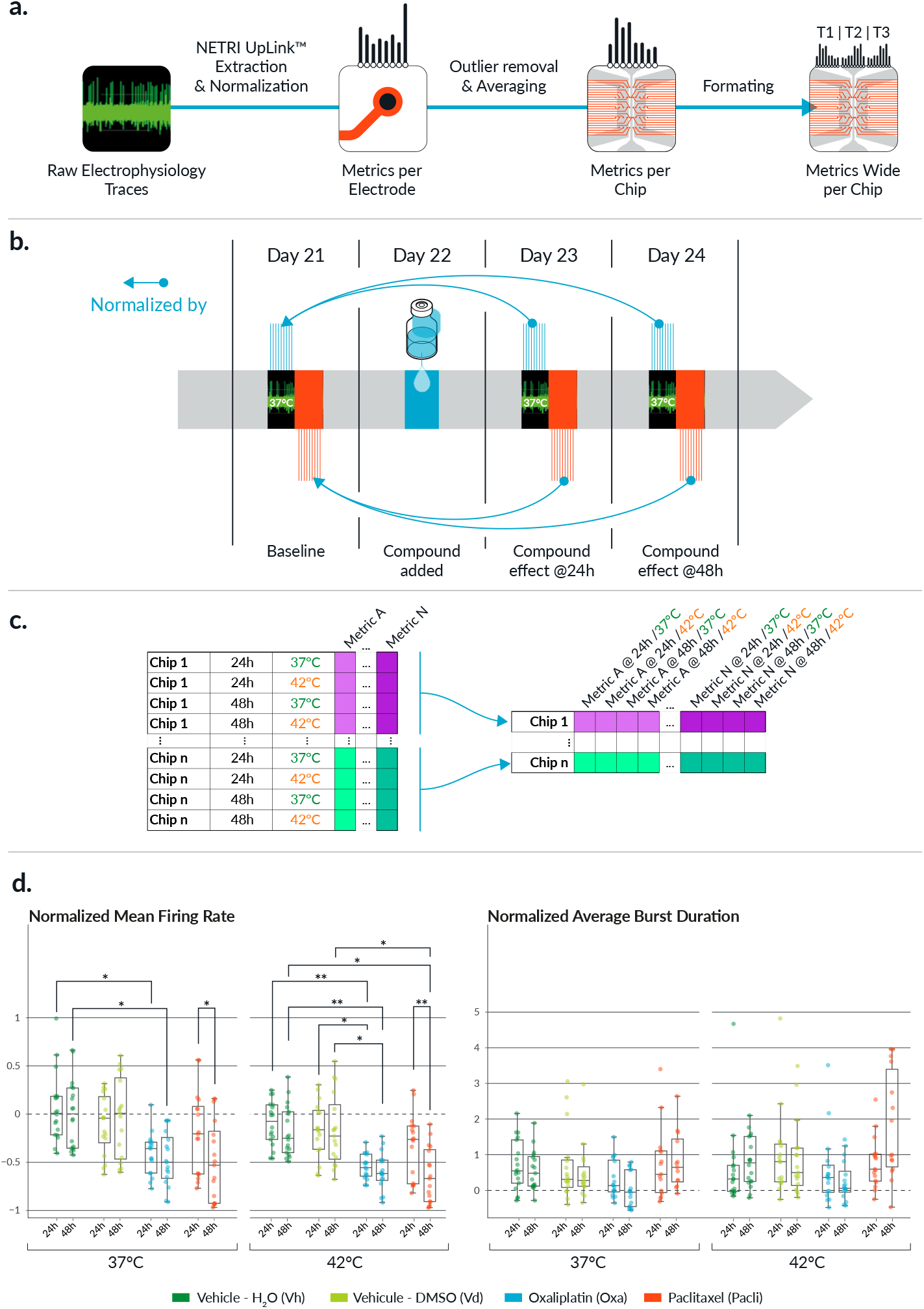
Data Acquisition and Preprocessing Pipeline. **a**. Schematic overview of the data preprocessing workflow. **b**. Normalization procedure: Each electrophysiological metric is normalized relative to its baseline value at the same temperature (pre-treatment, t_0_). **c**. Data formatting step: The dataset is transformed from long to wide format, with each row representing all features for a single MEA chip. **d**. Normalized Mean Firing Rate (left) and Normalized Average Burst Duration (right) for different compounds across time points and temperatures. Box plots show median (horizontal line) and interquartile range (boxes). *Statistical test:* Repeated-measures ANOVA with FDR correction. *p < 0.01, **p < 0.001.

The next step of the methodology consists of removing outliers before averaging the metrics at the chip level (see **Materials and Methods – Preprocessing**). This results in a data frame containing all the metrics associated with each recording at each different time point and temperature in a “long” format. It is then re-formatted in a “wide” format, such that each line in the data frame regroups all the metrics associated with any given chip (**Figure 3.c**). Each chip is thus associated with the same metrics at different timepoints or stimulation, describing its basal state and evolutive excitability.

This data formatting pipeline was applied to chips treated with oxaliplatin, paclitaxel, and their respective vehicles. Two commonly used MEA metrics, Mean Firing Rate (MFR) and Average Burst Duration (ABD), are shown in Figure 3d for different treatments, time points and temperatures. For the normalized MFR, significant reductions were observed for both oxaliplatin and paclitaxel compared to their respective vehicles at both 37°C and 42°C, particularly at the 48-hour timepoint. These results are consistent with previously reported effects of these compounds on neuronal activity^21^. Notably, the normalized MFR significantly decreases at 24h post-treatment for oxaliplatin while the effect of paclitaxel became significant only at 48 hours, with a more pronounced reduction observed at 42°C. In contrast, normalized ABD did not show significant differences between treatment groups. This indicates that while normalized MFR is a sensitive marker of compound effects in this context, normalized ABD alone may not provide sufficient discriminative power.

These findings highlight the differential effects of the two reference compounds and support the relevance of the NaaS methodology in modeling CIPN-related functional alterations. However, interpreting electrophysiological responses is inherently complex, as it requires accounting for both temporal dynamics and stimulus-dependent changes (e.g., temperature) across a wide array of metrics. To address this complexity, a dimensionality reduction approach is employed, enabling the identification of discriminative axes within the high-dimensional dataset and facilitating the visualization and classification of compound-specific effects.

### The NaaS discrimination Map

To obtain the map supporting the NaaS methodology, the data resulting from the preprocessing detailed in the previous section were fed into a dimensionality reduction model. Principal Component Analysis (PCA) was selected for its straightforward implementation and convenience of interpretation. The pair of principal components (PC) providing the highest discrimination were selected for the NaaS model (**Figure 4.a**).

**Figure 4.**
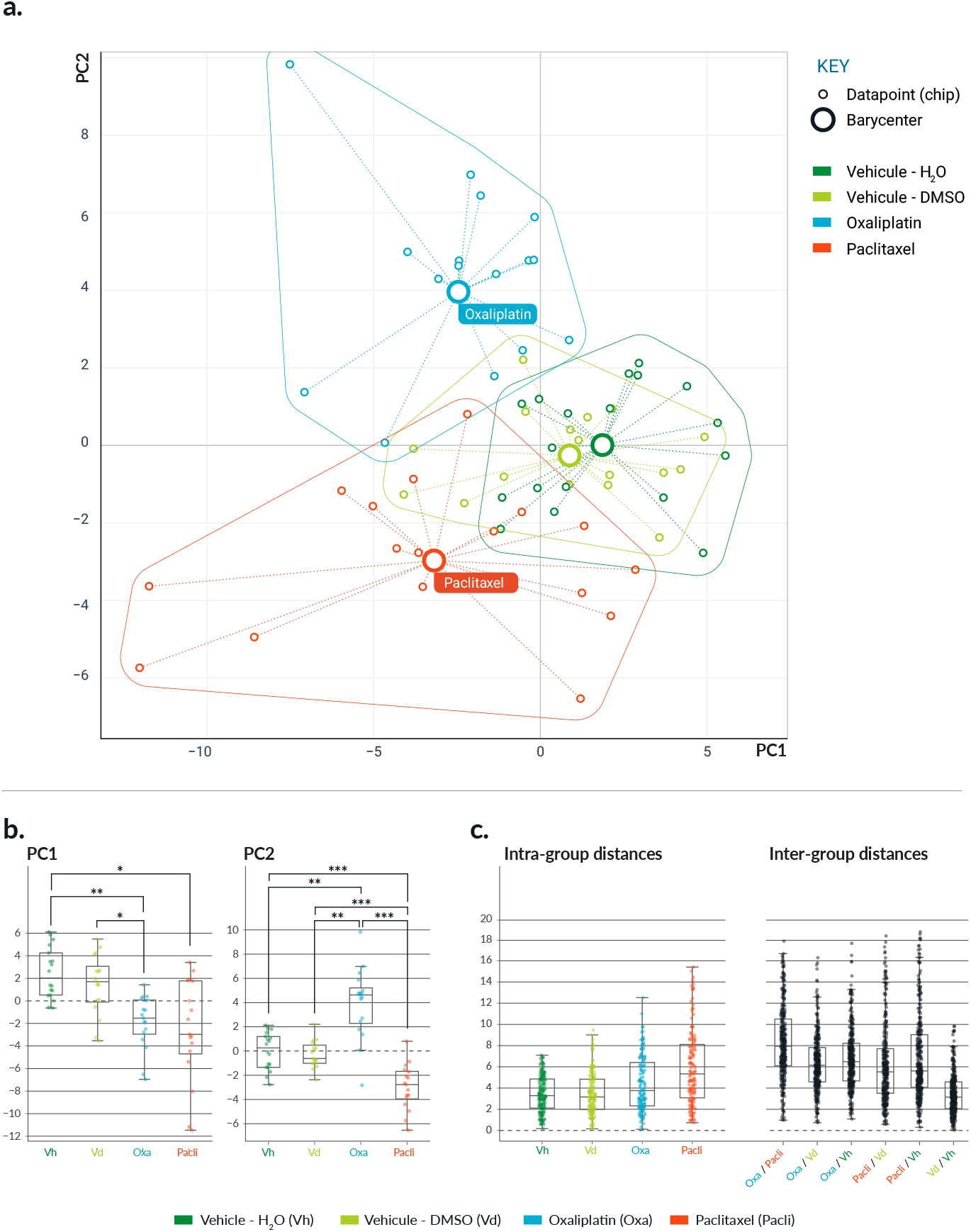
Functional Discrimination of Compounds in the NaaS Map. **a**. Projection of data points onto the first two principal components (PC1 and PC2), which explain 19.2% and 12.4% of the total variance, respectively. Solid circles represent group barycenters; plain lines depict group envelopes. **b**. Distribution of data points along PC1 (left) and PC2 (right). **c**. *Left:* Intra-group distances between replicates of the same compound. Each dot represents a distance between two chips. *Right:* Inter-group distances in the PC1–PC2 space, computed between each pair of compounds. *Statistical test:* Repeated-measures ANOVA with FDR correction. *p < 0.01, **p < 0.001, ***p < 0.0001.

In this map, the selected compounds are qualitatively well separated from each other as well as from their associated vehicle. This was quantitatively confirmed when examining the values alongside each PC, which are presented in **Figure 4.b**. Along PC1, both oxaliplatin and paclitaxel are statistically different from the vehicles (p<0.01), but no significant difference was observed between the two compounds. On the other hand, values along PC2 exhibited clear statistical differences between oxaliplatin and paclitaxel (p<0.001) and between the compounds and the vehicles (p<0.0001). Importantly, both vehicles displayed similar and consistent values for both PC.

The separability of the different compounds was further assessed by quantifying the intra and inter-group distances (**Figure 4.c**). The intra-group distance refers to the average distance between pairs of points from the same compound. As for the inter-group distance, it was defined for groups A and B as the average distance between each point in group A to all points in group B. The intra-group distances of both vehicles are comparable and close to their inter-group distance, thus providing an estimate of the inherent variability of the platform. This variability is lower than the intra-group distance of paclitaxel, whose neurotoxic effect likely induces a higher dispersion. With regards to the inter-group distances, they were similar for oxaliplatin and paclitaxel with respect to the vehicles, while the distances oxaliplatin / paclitaxel and vehicles DMSO / H20 were respectively higher and lower (**1**). These values highlight the separability of the different compounds when compared to the inter-group distance between vehicles, thereby confirming the choice of the subspace.

## DISCUSSION

In this paper, we proposed the NaaS methodology, consisting of a defined biological system, a micro-physiological device, a structured functional assay and a tailored analysis pipeline. The goal of the NaaS approach is to characterize and predict compound-induced functional changes in neurons by comparing their electrophysiological profiles to those of well-characterized reference compounds. By creating a multidimensional “map” of neuronal responses, the platform enables the positioning of unknown compounds within a reference space. This strategy can also be extended to evaluate the potential of companion therapies to mitigate adverse neurological effects caused by chemotherapeutic agents.

We demonstrated the feasibility of building a discriminating subspace supporting the NaaS methodology in the context of CIPN. Using a co-culture of hiPSC-derived sensory neurons and astrocytes seeded into compartmentalized MEA devices, we obtained robust electrophysiological data in response to two clinically relevant CIPN-inducing agents: paclitaxel and oxaliplatin. Our analysis showed a clear separation between the electrophysiological signatures of these two compounds, as well as from vehicle controls, confirming the platform’s potential to discriminate between distinct functional neurotoxic phenotypes. To further improve the map obtained, we foresee two main complementary paths: increasing separability (i.e., the distance between compounds), and lowering the variability (i.e., the dispersion within chips of the same compound).

From an experimental design perspective, a deeper and broader functional characterization of sensory neuron excitability could improve both separability and relevance. In this study, we focused on thermal hyperalgesia by implementing a heat stimulation at 42°C. However, other clinically relevant symptoms such as cold allodynia, particularly associated with oxaliplatin exposure, were not tested. To address this, future experimental designs could incorporate cold stimulation protocols or pharmacological TRP channel challenges in the terminal ending channel. For example, allyl isothiocyanate (AITC) for TRPA1 channels activation have been shown to model cold sensitivity in rodent DRG neurons ^22^. Including such stimulations would help isolate drug-specific phenotypes and enrich the electrophysiological dataset with new dimensions of sensory dysfunction.

From the biological system perspective, increasing the system’s complexity could enable the capture of additional pathophysiological mechanisms relevant to CIPN. One promising direction involves incorporating dorsal horn neurons, which play a crucial role in the processing and integration of sensory signals in the spinal cord. The functional synapse between peripheral sensory neurons and dorsal horn interneurons has been implicated in the development of central sensitization and chronic pain syndromes, including hyperalgesia ^23^.

Incorporating dorsal horn neurons into the experimental setup, using the TriaLink MEA device from NETRI for a compartmentalized culture, could enrich the study with network-level interactions between neuronal subtypes. Dorsal horn neurons also exhibit synchronized bursting activity, and changes in this network behavior could serve as an additional set of features to define compound-induced disruptions.

In terms of data processing, the most direct way of improving both separability and variability is to improve the data extraction and metrics calculations. In particular, it has been shown that the method selected for burst detection can still be optimized^24^. It is especially important in our context as more than two thirds of the metrics used depend on the burst detection. Furthermore, there is clear evidence that a substantial number of electrodes comprise signals arising from multiple units, and we would therefore highly benefit from spike sorting the data before the metrics calculation.

Moreover, to fully enable compound screening, the platform requires a broader panel of reference compounds for comparison and prediction of diverse adverse effects. Additional representatives of taxanes and platinum-based agents should be tested to determine whether they induce similar electrophysiological alterations. Other relevant classes to include are Vinca alkaloids, such as vincristine, as well as emerging therapies like antibody–drug conjugates (ADCs) and their payloads (e.g., MMAE).

Finally, the NaaS methodology is not conceptually limited to CIPN. It could be applied to other PNS domains such as inflammatory response or emetic response for gastrointestinal toxicity. Furthermore, in our platform, the nerve endings are isolated from their somas and can be coaxed to innervate another cell culture, be it 2D or 3D, representing a target organ. It paves the way towards other fields including dermatology with keratinocytes or 3D skin models, and toxicity evaluation with cardiac cells or liver organoids. Last, the methodology can also be adapted to central nervous system (CNS) domains such as neurotoxicity or seizure effect using CNS-specific neurons as sensors.

## MATERIALS AND METHODS

### MEA microfluidic devices, preparation and coating

DuaLink-MEA devices (NETRI, NF-DL-MEA) were left one hour under UV and one hour degassing in a vacuum chamber before use. Microfluidic channels were opened with 70% ethanol and immediately rinsed with PBS. Chips were then coated with 0.1mg/mL poly-D-lysin (PDL) (Thermofischer, A38904) overnight in a cell culture incubator. After 3 washes with PBS, channels were incubated for 2 hours with iMatrix 511-Silk (Anatomic, M511S) diluted 1:100 in the culture medium (SensoMM supplemented with Brainfast Astro Supplement diluted 1:1000). The iMatrix solution was removed prior to cell seeding.

### Cell culture and treatments

Sensory neurons were differentiated from human pluripotent stem cells obtained from CD34+ cord blood cells (Anatomic, RealDRG 1020F1-3M). Cortical astrocytes were differentiated from human hiPSC line (BrainXell, BX-0600).

40k sensory neurons and 20k astrocytes were seeded in the neuronal cell body compartment in 3µL of SensoMM (Anatomic, 1030) supplemented with Brainfast Astro Supplement (BrainXell, BX-2600) (1:1000). Brainfast Astro Supplement was maintained in the culture medium up until day 9. On day 1, the medium was fully renewed. Subsequently, 75% of the medium was replaced three times per week. Cells were cultured for 21 days before any assay.

Cells were treated at day 22 with Paclitaxel 1µM in 0.01% DMSO (Sigma, T7191) and oxaliplatin 10µM in 0.125% water (Cliniscience, NB-64-00307). Vehicles Vd and Vh correspond respectively to medium with these concentrations of DMSO or water.

### Immunostaining and image acquisition

Cells were fixed at day 24 with 4% formaldehyde (Thermofisher, 28908), permeabilized with Triton 0.1% (Sigma, X100) and blocked with 1% BSA (Thermofisher, 14190250).

Cells were stained with the following primary antibodies: β-III tubulin (Thermofisher, MA1-118), MAP2 (Abcam, ab5392) and GFAP (Abcam, ab16997) overnight at 4°C. Devices were incubated with secondary antibodies: donkey anti-mouse IgG (Abcam, ab150110), donkey anti-rabbit IgG (Abcam, ab150075), goat anti-chicken IgY (Abcam, ab150169) and goat anti-mouse IgG (Abcam, ab150113) 2 hours at room temperature. Nuclei were counterstained with 3 µM DAPI (Sigma, D8417).

Fluorescent images were acquired with an Axio Observer 7 microscope (Carl Zeiss).

### Microelectrode array recording

For each recording, plates were equilibrated for 15 minutes in the MaestroPro (Axion Biosystems). Recordings are made with 5% CO_2_ in spontaneous activity mode.

At day 21 plates were recorded for 5 minutes at 37°C and at 42°C. Activity from the 37°C recording has been used to allocate each device in homogeneous treatment groups. The next day, devices were treated with the different compounds or vehicles by a full medium change in the three channels.

After 24h of treatment, all chips were recorded for 5 minutes at 37°C and 5 minutes at 42°C followed by a 50% culture medium change containing treatments.

After 48h of treatment, all chips were recorded for 5 minutes at 37°C and 5 minutes at 42°C.

### Metrics calculation

The data extraction process was performed with the software AxisNavigator (Axion Biosystems, v3.12.6), including spike detection, burst detection and metrics calculation at the electrode level. Compartmentalization was managed with the NETRI UpLink Software (NETRI, v1.4.1), and only electrodes in the micro-channels were kept to maintain a high signal-to-noise ratio.

The spikes were identified with AxisNavigator with default parameters (adaptive threshold correction, threshold 6xSTD). Then, burst detection was performed using the Poisson Surprise method^25^, with a surprise threshold of 10.

The following list of metrics was computed:

|  |  |  |
| --- | --- | --- |
| Number of Spikes | Mean Firing Rate (Hz) | ISI Coefficient of Variation |
| Number of Bursts | Burst Duration Avg (s) | Burst Duration Std (s) |
| Number of Spikes per Burst Avg | Number of Spikes per Burst Std | Mean ISI within Burst Avg |
| Mean ISI within Burst Std | Median ISI within Burst Avg | Median ISI within Burst Std |
| Inter Burst Interval Avg (s) | Inter Burst Interval Std (s) | Burst Frequency (Hz) |
| IBI Coefficient of Variation | Normalized Duration IQR | Burst Percentage |

### Statistics

All statistical comparisons presented were performed using pairwise tests corrected for multiple comparisons using the Pingouin (v0.5.5) Python library.

### Preprocessing

Once metrics were computed at the electrode level, they were normalized as a relative change with respect to the basal recording acquired at the same temperature. Outliers were detected by computing the Z-score with respect to the distribution within the chip, and a threshold of 9 was applied to remove the electrodes with extreme values. The remaining values are then averaged to obtain the metric at the chip level. The resulting data frame was converted from the “long” to the “wide” format, such that each line in the frame comprises all metrics at all temperatures and time points associated with one chip. All operations were developed in Python 3.12 using the “pandas” library (v2.2.3).

### NaaS map

To obtain the map supporting the NaaS model, dimensionality reduction was performed on the dataset with Principal Component Analysis. The two first axes were kept as retaining the most variance. The implementation was done using scikit-learn (v1.6.1).

## ACKNOWLEDGEMENT

We thank Hélène Gauthier and Rania Talbi for their scientific contributions in shaping the project and laying its foundations. We are grateful to Benoît Maisonneuve and Adriana Toma for their valuable advice throughout the work. We also acknowledge Anatomic Inc., and in particular Vince Truong, for insightful discussions and support in improving culture quality.

